# Modulation of *Saccharomyces cerevisiae* Stm1_N^1-113^and human Aβ42 amyloid fibril morphology by 3,3’-(acridine-4,5-diylbis(methylene))bis(1-(carboxymethyl)-1H-benzimidazolium) dibromide

**DOI:** 10.1101/2025.08.04.668593

**Authors:** Patil Pranita Uttamrao, Uttam Das, Rohith R Kumar, Dinesh Harijan, Ashwath Kumar Balu, D Krishna Rao, Ganesan Prabusankar, Thenmalarchelvi Rathinavelan

**Affiliations:** Department of Biotechnology, Indian Institute of Technology Hyderabad, Telangana State- 502284, India; Department of Chemistry, Indian Institute of Technology Hyderabad, Telangana State- 502284, India; Tata institute of fundamental research Hyderabad, Telangana State- 500046, India

**Keywords:** Stm1, amyloid, *Saccharomyces cerevisiae*, acridine derivative, AFM, NMR, Aβ42

## Abstract

Acridine derivatives are among the oldest and most effective classes of chemotherapeutic agents, exhibiting a broader spectrum of bioactivities like antitumor, antifungal, antimicrobial, and antiviral. They remain a preferred choice for molecular imaging of amyloid as well as for inhibiting amyloid fibrillation or stabilizing the fibril. This study reports the effect of a newly reported acridine compound 3,3’-(acridine-4,5-diylbis(methylene)) bis(1-(carboxymethyl)-benzimidazolium) dibromide (henceforth, Ac-BIM-acid) on the amyloid-like fibrillation of N-terminal region of *Saccharomyces cerevisiae* Stm1 protein (Stm1_N^1-113^) and human Aβ42. Modulation of Stm1_N^1-113^ amyloid-like structures at 400 µM concentration with respect to varying concentrations of Ac-BIM-acid is revealed by AFM and NMR. While 2D-HSQC NMR spectra show the binding of Ac-BIM-acid with 400 µM Stm1_N^1-113^, AFM captures the morphological changes of Stm1_N^1-113^ in response to 1 mM and 2.5 mM Ac-BIM-acid at physiological salt concentration in a time-dependent manner. Similarly, Ac-BIM-acid is shown to modulate the amyloid morphology of human Aβ42 protein, responsible for Alzheimer’s disease, captured in AFM. Docking studies carried out indicate that the hydrophobic acridine ring binds at the hydrophobic pocket of the N-terminal β-sheet and it’s neighboring β-sheet of the Aβ42 monomorphic fibril. Thus, Ac-BIM-acid would be yet another addition to the class of acridine derivatives that modulate amyloid fibrillation.

## 1. Introduction

Acridine derivatives are among the preferred bioactive compounds that can bind to amyloid structures, which are associated with various neurodegenerative diseases such as Alzheimer’s, Parkinson’s, and Huntington’s disease (Grayson et al., 2021). Amyloids follow a hierarchical structural organization: monomer, oligomer, protofibril, and fibril. Amyloid formation occurs due to the misfolding or alteration(s) in protein conformation, resulting in beta-strand formation (monomer). These monomers start stacking onto each other, forming oligomeric to long fibrils (Berthelot et al., 2013). These amyloid structures deposit in the form of insoluble extracellular/intracellular plaques, causing detrimental effects (Berthelot et al., 2013).

Several ligands have been identified to bind with amyloid structures (Chen et al., 2017; Chisholm et al., 2024; Chisholm et al., 2023; LeVine et al., 2016; Salahuddin et al., 2021; Sharma et al., 2023; Yao et al., 2022; Young et al., 2017). These are utilized for various applications ranging from primary research and diagnostics to treatment. Among them, acridine derivatives have drawn significant attention due to their unique structural properties and diverse therapeutic applications (Antosova et al., 2011; Gabriel, 2020; Makhaeva et al., 2023; Monika Gensicka-Kowalewska, 2017; Prasad et al., 2016; Sharma et al., 2023; Varakumar et al., 2022). The planar and tricyclic structure of the acridine moiety facilitates the intercalation and stacking interactions with the β-sheet-rich regions of amyloid fibrils (Antosova et al., 2011). The acridine derivatives have diverse applications, which include the inhibition of amyloid aggregation (Basagni et al., 2022), disaggregation of existing fibrils (Low et al., 2022; Noi et al., 2021), and stabilization of non-toxic intermediate forms. They have also been used as imaging agents to detect the amyloid plaques in the tissues of patients affected by neurodegenerative disorders (Leuzy et al., 2019; Mason et al., 2013; Mathis et al., 2012; Saint-Aubert et al., 2017). However, there is no drug molecule of any kind available in the market to cure any of the amyloid diseases. Thus, developing new acridine derivatives is essential for advancing our understanding of amyloid-related disorders and their treatment options.

Yeast models are widely used to study amyloid diseases and their therapeutics (Rencus-Lazar et al., 2019). Recently, the N-terminal region of *Saccharomyces cerevisiae* Stm1 (suppressor of the target of Myb protein 1) (Stm1_N^1-113^) has been shown to form amyloid-like conformation with increasing concentration of protein (100 to 400 µM) (Subbaiah et al., 2023). It is noteworthy that although we have reported earlier that Stm1_N1-113 forms amyloid conformation (Subbaiah et al., 2023), it is being mentioned in this study as an “amyloid-like” conformation as the Stm1_N1-113 fibril dimensions are different from the conventional amyloid fibril dimension. The propensity to form amyloid-like fibril is also shown to increase with time (0, 24, and 48 h). Stm1 protein is a multifunctional protein involved in regulating ribosome homeostasis under stress conditions such as nutrient deprivation and oxidative stress (Koli et al., 2024). The accumulation of Stm1 protein has been demonstrated to cause cell death in yeast (Ligr et al., 2001). It was proposed earlier that amyloid-like formation is the molecular basis for such cell death (Subbaiah et al., 2023). Thus, the amyloidogenic characteristic of Stm1_N^1-113^ is exploited here to study the influence of a recently reported acridine derivative (Revi et al., 2025) on the amyloid-like conformation. Atomic force microscopy (AFM) and heteronuclear single quantum coherence (HSQC) NMR are employed to investigate the ability of 3,3’-(acridine-4,5-diylbis(methylene))bis(1-(carboxymethyl)-1H-benzimidazolium) dibromide (henceforth, Ac-BIM-acid) to bind with Stm1_N^1-113^. This is further extended to investigate the effect of Ac-BIM-acid on amyloid fibrillation of human Aβ42 using AFM and docking studies.

## 2. Materials and methods

### 2.1 Ac-BIM-acid synthesis and preparation

3,3’-(acridine-4,5-diylbis(methylene))bis(1-(carboxymethyl)-benzimidazolium) dibromide (Ac-BIM-acid) was synthesized using the previously reported procedure with minor modifications (Gembali Raju, 2016; Prasad et al., 2016). The reactions were carried out under an argon atmosphere. The solvents were distilled under an argon atmosphere. The multi-nuclear magnetic resonance spectra of Ac-BIM-acid were collected using Bruker Ultra shield 400 MHz spectrometers at 25 °C to confirm the synthesis. Fourier transform infrared (FT-IR) data were collected in a Bruker Alpha-P Fourier transform spectrometer for the compound to further validate Ac-BIM-acid synthesis. A stock solution of 10 mM Ac-BIM-acid was prepared by solubilizing it in milliQ water and filtered using a 0.2 µm syringe filter.

### 2.2 Sub-cloning, expression, and purification of N-terminal domain of *Saccharomyces cerevisiae* Stm1

The gene sequence corresponding to Stm1_N^1-113^ was sub-cloned into a pDZ1 expression vector (a modified pET-21A vector with a T7 promoter and ampicillin-resistance gene), as mentioned elsewhere (Subbaiah et al., 2023). The 5’end of the gene (*viz*., the N-terminus of the protein) is immediately flanked by a Tev protease site, which connects the His_5_ and GB1-tags to the gene (Rathinavelan et al., 2014; Rathinavelan et al., 2011). The pDZ1 vector coding for Stm1_N^1-113^ was transferred into *E. coli* BL21 (DE3) cells. To overexpress the protein, a 10 mL pre-inoculum culture was grown overnight and transferred into 1 L Luria-Bertan (LB) broth with 100 mg of ampicillin. The culture was grown at 37 °C and 210 rpm until the OD reached 0.6. Subsequently, the culture was induced with 1 mM isopropyl β-d-1-thiogalactopyranoside (IPTG) and incubated at 25 °C overnight. The harvested cells were sonicated in binding buffer (20 mM Tris-HCl, 500 mM NaCl, 5 mM imidazole, pH 8.0) with 600 µl of 0.1 mM for 30 ml harvested cells) phenylmethanesulfonyl fluoride (PMSF) to inhibit protease activity. After centrifugation at 13000 rpm for 10 min, the supernatant was added with 5% polyethyleneimine (PEI) (300 µl) and centrifuged again at 13000 rpm for 10 min. The supernatant obtained after the final centrifugation was subjected to purification. The purification encompasses two steps in Ni^2+^-NTA affinity column chromatography. During the first round of purification, the protein attached to the tags was collected in the elusion buffer (20 mM Tris-HCl, 500 mM NaCl, 200 mM imidazole, pH 8.0). In the next step, the His6 and GB1 tags at the N-terminus of Stm1_N^1-113^ were cleaved using TEV protease in an overnight dialysis (20 mM Tris-HCl buffer (pH-8.0), 0.5 mM EDTA, 2 mM DTT, 20 mM NaCl). During the next round of purification, Stm1_N^1-113^ cleaved from the tags was collected in the binding buffer, dialyzed in phosphate buffer (10 mM sodium phosphate with 150 mM NaCl, pH 7.4), and concentrated using an Amicon protein concentrator. Protein concentration was measured at 280 nm using a UV spectrophotometer with an extinction coefficient of 5500 (mg/ml)^-1^cm^-1^. For the biophysical studies other than NMR, the protein was expressed and purified as mentioned above.

For the synthesis of ^15^N labelled Stm1_N^1-113^ protein to use in NMR experiments, *E. coli* BL21 (DE3) cells carrying the plasmid were cultured in 1 L M9 minimal medium by using the pre-inoculum culture grown overnight in LB media. The M9 minimal medium contained 3 g of glucose and 0.2 g of ^15^N-ammonium chloride per liter, along with essential trace metals and vitamins, as previously reported (Rathinavelan et al., 2011; Wang et al., 2007). The rest of the protein expression and purification steps were the same as described above. Exceptionally, 50 mM NaCl was used instead of 150 mM during the final dialysis to facilitate NMR experiments.

### 2.3 Preparation of Aβ42 amyloid fibrils

Lyophilized Aβ42 peptide was purchased from Biotech Desk Pvt Ltd. The vial containing the peptide was initially subjected to centrifugation at 3000rpm at room temperature. Subsequently, a 200 µM stock solution was prepared by dissolving the peptide in double-deionized water. The solution was vortexed gently to ensure complete dissolution. To avoid degradation, the solution was immediately aliquoted and stored at -20 °C until use. Thawed aliquots were used for the experiments. A 100 µM concentration of the peptide (at 0, 24, and 48 h) was used in the circular dichroism (CD) and AFM experiments.

### 2.4 Atomic force microscopy

AFM images of Stm1_N^1-113^ and Aβ42 amyloid morphologies were captured using Park Systems NX10 in tapping mode, and the images were analysed using XEI software. 400 µM of Stm1_N^1-113^ protein in phosphate buffer and 150 mM NaCl was titrated with 1 mM and 2.5 mM Ac-BIM-acid and incubated for 30 mins (0 h), 24 h and 48 h at 25° C. Similarly, 100 µM Aβ42 was incubated with 5 mM Ac-BIM-acid for 0, 24 and 48 h. Following this, Stm1_N^1-113^ and Aβ42 were independently deposited on an APTES-coated mica sheet (Shlyakhtenko et al., 2013) to facilitate AFM data collection. The mica sheets were then washed and dried following a standardized protocol ensuring the removal of excess salts (Shlyakhtenko et al., 2013). Using the same protocol, the time-dependent AFM images of Stm1_N^1-113^ and Aβ42 were captured independently. Detailed methodology can be seen in a previous study (Subbaiah et al., 2023).

### 2.5 Two-dimensional NMR experiments

Two-dimensional ^1^H-^15^N HSQC spectra corresponding to 400 µM of ^15^N-labelled Stm1_N^1-113^ protein titrated with 0 mM and 0.5 mM concentrations of Ac-BIM-acid (incubated for 1 hr at 25 °C were recorded. Buffer conditions were maintained at pH 7.4 with 10 mM sodium phosphate buffer, 50 mM NaCl, and 10% D_2_O. All the NMR experiments were carried out using a gradient-based HSQC pulse program along with solvent suppression at 25 °C on a 500 MHz Bruker NMR spectrometer with a 5 mm triple resonance inverse probe (TXI probe) by maintaining the same acquisition parameters for all the experiments. NMR Spectra were recorded with 32 scans in the direct and 150 points in the indirect dimension for 0, 24, and 48 hours of incubation of Ac-BIM-acid with Stm1_N^1-113^. NMR data was processed using Topspin

4.2.0 version software and NMR ViewJ (Johnson, 2004). The exact contour threshold level was maintained for all HSQC spectra reported in this study to obtain the peak intensity. The peak intensity was obtained after peak picking in the topspin, and the peak intensity was taken from the list of prominent HSQC signals.

GraphPad Prism 8.0.2 software was utilized for making bar plots in AFM and NMR data analysis (Windows).

### 2.6 Docking of Ac-BIM-acid with Aβ42 amyloid

Docking of Ac-BIM-acid with Aβ42 monomorphic amyloid fibril structure (receptor) (**PDB ID: 2MXU**) solved by solid state NMR (Xiao et al., 2015) was carried out using AutoDock 4.2 (Morris et al., 2009) uses genetic algorithm (GA) to search the ligand binding site to a receptor. Initially, the ligand was converted from SDF to PDB format using Open Babel (O’Boyle et al., 2011). Prior to docking, the ligand (Ac-BIM-acid) and receptor (Aβ42 fibril) were added with the hydrogen atoms and partial charges using AutoDock tools (ADT). 10 (suggested to be ideal) torsional degrees of freedom (*viz*., rotation around single bonds within a molecule) was used to incorporate the flexibility to the ligand and exhaustiveness (*viz*., number of reputations of initial random runs) value of 8 (default) was used for search and optimize the conformational space of ligand. The receptor was kept rigid during the docking. A grid size of 90x90x90 Å, 100 independent GA runs, and 250,000 energy evaluations per genetic algorithm run (viz., 250,000*100 runs = 250,00,000) were considered for the blind docking simulation. Gasteiger partial atomic charges were considered to calculate the electrostatic interaction between the receptor and ligand. The lowest binding free energy conformation among the docked structure was considered for visualization and analysis using PyMol and Discovery studio 2025 respectively.

## 3. Results

### 3.1 Characterization of 3,3’-(acridine-4,5-diylbis(methylene))bis(1-(carboxymethyl)-benzimidazolium) dibromide (Ac-BIM-acid)

3,3’-(acridine-4,5-diylbis(methylene))bis(1-(carboxymethyl)-1H-benzimidazolium) dibromide (Ac-BIM-Acid) (**Figure 1**) was synthesized with an excellent yield of 83% by following the previously reported procedure (Gembali Raju, 2016 ; Prasadetal., 2016) . The formation of Ac-BIM-acid was confirmed by multinuclear 1D (^1^H and ^13^C) NMR (**Figures S1 and S2**) and FT-IR spectra (**Figure S3**).

**Figure 1.**
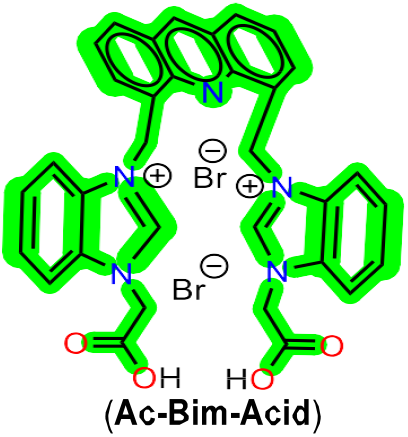
Schematic representation of Ac-BIM-Acid.

### 3.2 Titration of Ac-BIM-acid with Stm1_N^1-113^ amyloid

#### 3.2.1 Perturbation in the ^1^H-^15^N HSQC of Stm1_N^1-113^ upon titration with Ac-BIM-acid

To assess the binding of Ac-BIM-acid with 400 μM Stm1_N^1-113, 1^H-^15^N HSQC data is collected by titrating 0.5 mM Ac-BIM-acid at three different time intervals (0 h, 24 h and 48 h). Note that 400 μM concentration of Stm1_N^1-113^ is chosen as it was reported earlier that the protein forms proto-fibrils gradually with respect to time at this concentration (Subbaiah et al., 2023). In the presence of Ac-BIM-acid, chemical shift perturbations are seen in the HSQC with respect to time (**Figure 2A-C**). The perturbations are either due to the emergence of new peaks or due to the increase in the peak intensity at 48 h compared to 0 h and 24 h spectra (**Figure 2D&E**). Thus, the interaction between Stm1_N^1-113^ and Ac-BIM-acid becomes apparent with respect to time (**Figure 2D**).

**Figure 2.**
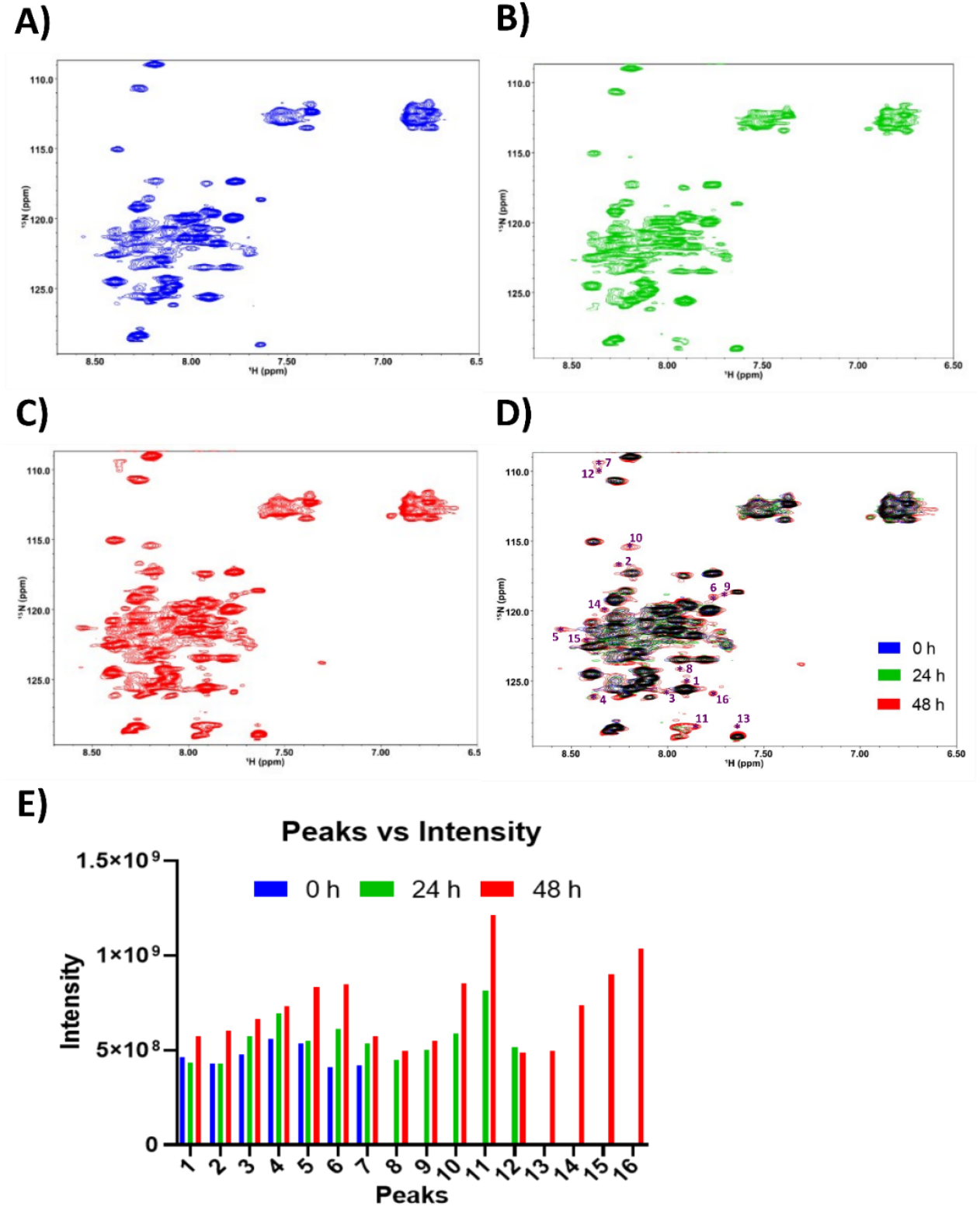
^1^H-^15^N HSQC spectra of 400 μM Stm1_N^1-113^ in the presence of 0.5 mM Ac-BIM-acid at A) 0 h, B) 24 h and C) 48 h and their (D) overlay. (E) Intensity comparison of peaks (marked by numbers 1 to 16 in (D)) that have shown perturbation with respect to time.

At 24 h, 5 new peaks have emerged (marked by 8 to 12), which are absent at 0 h (**Figure 2E**). Further, there is an increase in the peak intensity (compared to 0 h) of already existing peaks (peaks 1 to 7) (**Figure 2E)**. These peaks continue the increasing trend at 48 h, too (peaks 1 to 12). More importantly, 4 new peaks have appeared at 48 h (peaks 13 to 16). While these HSQC spectra indicate the binding between Ac-BIM-acid with the Stm1_N^1-113^, they do not provide any information about the morphological alterations in Stm1_N^1-113^ upon binding with the ligand. This has led us to utilize the AFM technique to capture the alterations in the fibril morphology, if any.

#### 3.2.2 Stm1_N^1-113^ undergoes morphological changes upon binding with the Ac-BIM-acid

Atomic force microscopy (AFM) images of 400 μM Stm1_N^1-113^ in the absence and presence of 1 mM and 2.5 mM concentration of Ac-BIM-acid are captured at 0 h, 24 h, and 48 h. In the absence of Ac-BIM-acid at 0 h, the protein showed fibril formation (**Figures 3A and S4A**) with dimensions ranging from 0.7 to 1.4 µm in length and 110 to 200 nm in height. The binding of fibrils with 1 mM Ac-BIM-acid is seen to have minimal effect on the morphology of fibrils (with length ranging from 0.5 to 0.8 µm and height ranging from 110 to 300 nm) at 0 h (**Figures 3(B&D) and S4(B&D)**). In the presence of 2.5 mM Ac-BIM-acid, the change in the fibril dimensions starts becoming more apparent with length ranging from 0.4 to 0.6 µm and height ranging from 32 to 99 nm (**Figures 3(C&D) and S4(C & D)**).

**Figure 3.**
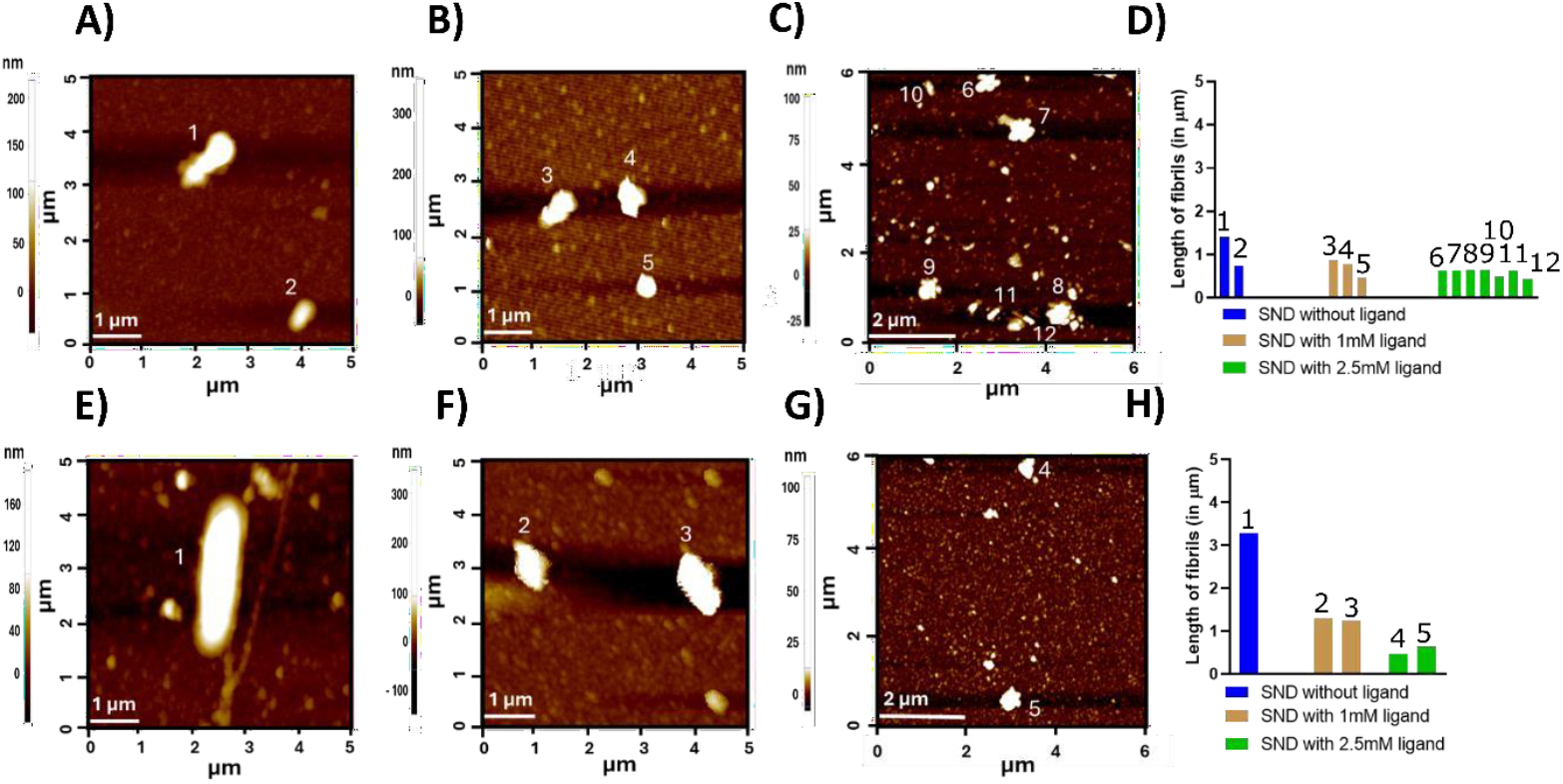
AFM images of 400 μM Stm1_N^1-113^ at (A, B & C) 0 h and (E, F & G) 24 h in the absence (A and E) and presence of 1 mM (B & F) and 2.5 mM (C & G) Ac-BIM-acid. Note that each fibril is marked with a number. The bar plots represent the comparative length and height of the fibrils in the presence (1 mM (golden color) and 2.5 mM (green color)) and absence (blue color) of Ac-BIM-acid at D) 0 h and H) 24 h.

At 24 hours, the presence of Ac-BIM-acid influences its morphology, leading to a noticeable shortening of the fibril length. 1 mM Ac-BIM-acid reduces the Stm1_N^1-113^ fibril length from 3 µm to 1.2 µm, which is almost half of the protein fibril length (**Figures 3(E, F &H) and S4(E, F &H)**). In the presence of 2.5 mM Ac-BIM-acid, the dimension of the fibril further gets reduced with a length of ∼ 0.5 µm and height of ∼ 70 nm (**Figures 3(G&H) and S4(G&H)**).

Interestingly, at 48 h, amyloid-like species of two different morphologies are seen in the case of protein alone: ∼4.5 µm long branched fibril with an average height of 200 nm and ∼2.5 µm long cylindrical fibril (**Figures 4(A & D) and S5(A&D)**). In line with 24 h data, the fibril morphology is significantly affected at 48 h in the presence of 1 mM Ac-BIM-acid: individual short fibrils of ∼1 µm length as well as a long fibril resulting from the lengthwise merging of these short fibrils are observed (**Figures 4(B & D) and S5(B&D)**). However, 2.5 mM Ac-BIM-acid completely inhibits the fibrillation at 48 h, resulting in only spherical oligomeric structures (**Figures 4(C & D) and S5(C&D)**).

**Figure 4.**
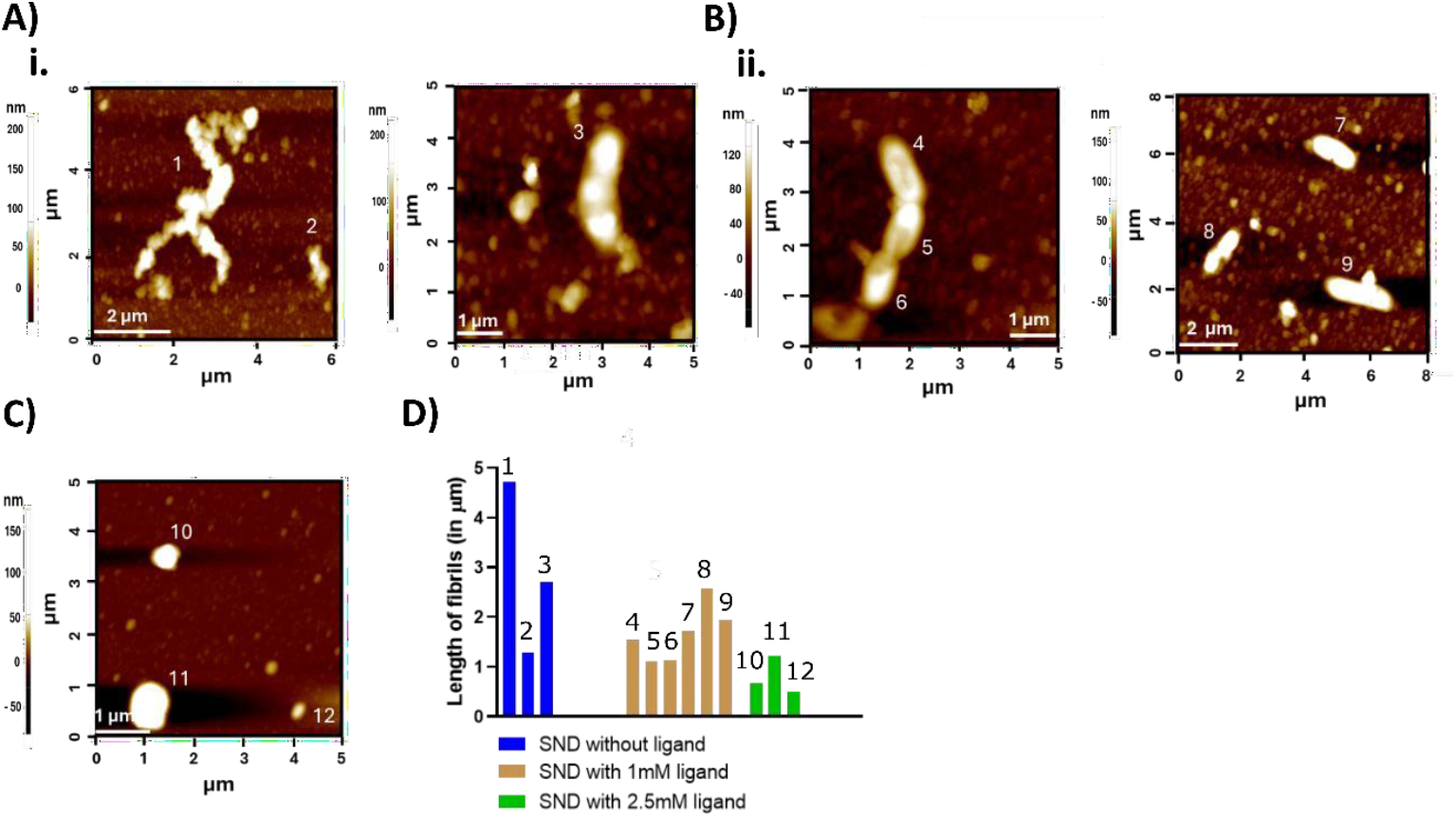
AFM images of 400 μM Stm1_N^1-113^ at 48 h in the absence (A) and presence of 1 mM (B) and 2.5 mM (C) Ac-BIM-acid. Note that each fibril is assigned a number as marked in the images. The bar plots represent the comparative length of the fibrils in the presence or absence of Ac-BIM-acid at 48 h (D).

The AFM image captured for 1 mM Ac-BIM-acid alone at physiological salt concentration showed just a few small granular structures, confirming that the fibrils belong to Stm1_N^1-113^ (**Figure S6**). The results from ^1^H-^15^N HSQC and AFM, thus, show the alteration in Stm1_N^1-113^ amyloid-like fibril morphology by Ac-BIM-acid.

### 3.3 Characterizing the interaction between Ac-BIM-acid with Aβ42 amyloid

After confirming the Aβ42 fibril formation after 12 h onwards (at 25 °C) (**Figure S7**), AFM experiments are carried out. The AFM images of Aβ42 amyloid collected at 0, 24 and 48 h confirm (**Figure 5, S8**) the amyloid fibrillation as reported in earlier studies (Zhaliazka et al., 2023). While oligomers are seen at 0 h (**Figure 5A**), more crowded fibrils are seen at 48 h (**Figure 5C**), with 24 h (**Figure 5B**) exhibiting the mixture of oligomers and fibrils. 5 mM Ac-BIM-acid with Aβ42 amyloid incubation with 0, 24 and 48 h indicate the effect (**Figure 5, S8**) of the compound from 0 h onwards. While the density of oligomers are reduced at 0 h (**Figure 5D**), shorter fibrils are seen at 24 h (**Figure 5E**). However, at 48 h, the fibrils are totally not seen (**Figure 5F**); instead, crowded oligomers are seen, indicating that the compound inhibits the fibril formation.

**Figure 5.**
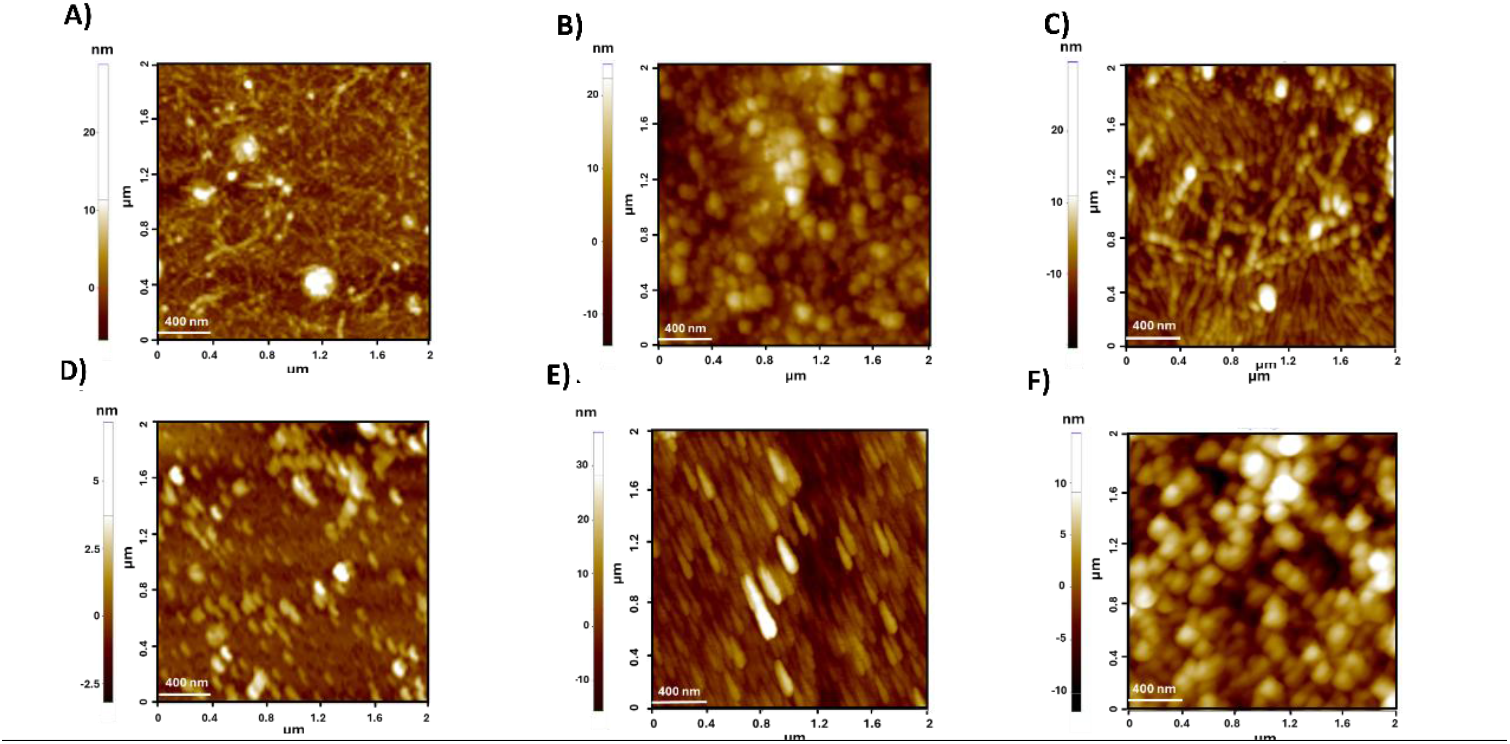
AFM images of 100 μM Aβ42 at (A & D) 0 h, (B and E) 24 h and (C & F) in the absence (A, B & C) and presence of 5 mM (D, E & F) Ac-BIM-acid.

### 3.4 Docking reveals Ac-BIM-acid sandwiches between A42 fibril β-sheets

Docking of Ac-BIM-acid with the Aβ42 amyloid fibril structure (PDB ID: 2MXU) indicates that they interact with each other favourably with the free energy of -9.75 kcal/mol, and they interact predominantly through van der Waal’s and hydrophobic interactions. Ac-BIM-acid binds in the hydrophobic cavity formed at the interface of Aβ42 N-terminal β-sheet and the other β-sheet in its immediate vicinity (**Figure 6A (Left)**). The cavity encompasses 12 stacked Aβ42 monomers. While the acridine ring of the ligand is aligned with N-terminal histidine-14 residues of 3 consecutive fibril monomers (**Figure 6A (Right)**), the two benzimidazole rings rest in the hydrophobic cavity formed by leucine-17, isoleucine-32 and glycine-33 (**Figure 6A (Right) and Figure 6B**). Notably, the carboxylic groups of 2 benzimidazole rings of the Ac-BIM-acid form a weak hydrogen bond (**Figure 6B**) with the backbone carbonyl group of glycine-33 belonging to two different monomers.

**Figure 6.**
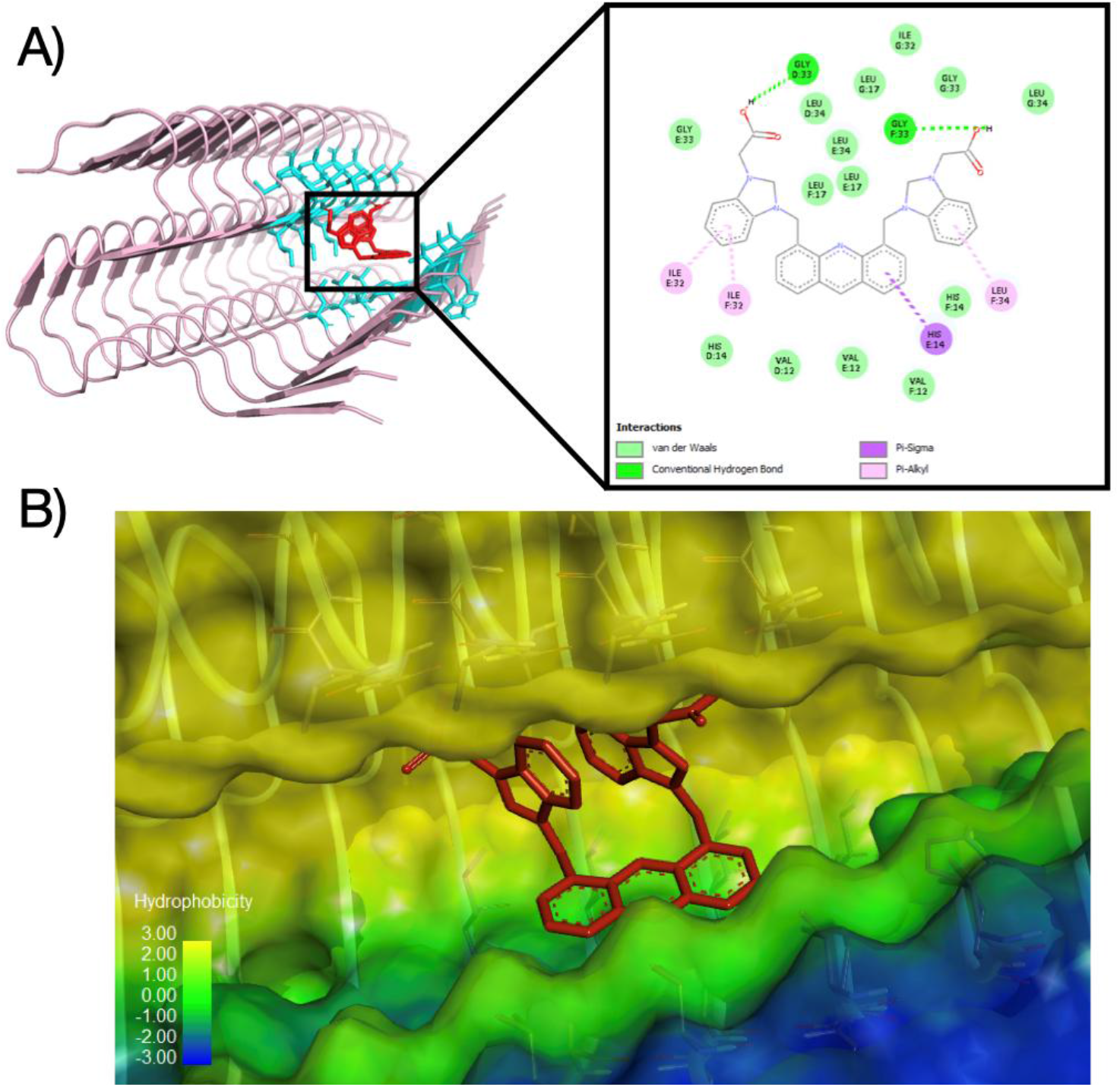
(A) Lowest energy docked complex of Ac-BIM-acid (red colored sticks) and Aβ42 fibril (cartoon representation) (Left). Note that His-14, Leu-17, ILE-32 and Gly-33 in the vicinity of the ligand are shown in cyan colored sticks in Left. Atomistic details about the interaction between Ac-BIM-acid and Aβ42 fibril is shown in the 2D-interaction map along with amino acids (color coded circles) involved in different non-bonded interactions (Right). Dotted lines indicate the amino acids that directly interact with ligand (Right). (B) Hydrophobicity profile at the binding pocket of the Aβ42 fibril is shown in surface representation. As can be seen, the interaction between them is Hydrophobically favourable (hydrophobicity=+3). Hydrophobicity scale is also shown for reference.

## 4. Discussion

Acridine derivatives are versatile bioactive compounds that are being widely utilized for various biomedical applications. Here, the binding interactions between a newly synthesized acridine derivative 3,3’-(acridine-4,5-diylbis(methylene))bis(1-(carboxymethyl)-benzimidazolium) dibromide (shortly, Ac-BIM-acid, **Figure 1**) and amyloidogenic Stm1_N^1-113^ protein is explored with a particular focus on the potential of Ac-BIM-acid in altering the morphology of Stm1_N^1-113^ amyloid-like structures. Earlier findings have shown that the N-terminal domain of Stm1 protein forms amyloid-like structures after a threshold concentration in a time-dependent manner (Subbaiah et al., 2023).

The ^1^H-^15^N HSQC-NMR and AFM results show the ability of Ac-BIM-acid to interact and modulate the amyloid-like fibrils of Stm1_N^1-113^ (**Figures 2-4 & S4-S5**). The appearance of new peaks and broadening of peaks seen in ^1^H-^15^N HSQC-NMR spectra in a time-dependent manner (0, 24, and 48 h) suggest local conformational changes in the protein upon binding with Ac-BIM-acid (**Figure 2**). AFM data of Stm1_N^1-113^ protein in the absence and presence of Ac-BIM-acid compound shows the effect of this interaction on the morphology of the amyloid-like fibrils in a time-dependent manner. It is noteworthy that Stm1_N^1-113^ protein fibrils observed here at 400 μM concentration in the absence of Ac-BIM-acid at 0, 24, and 48 h resemble the fibrils that were reported earlier (Subbaiah et al., 2023). Interestingly, the fibril species reported earlier at 48 h, formed through the height-wise stacking of the stable units (1 µm length), is not observed here. Instead, a branched and a single lengthy (2.7 μm length) species are seen (**Figure 4**). These species might have formed out of length-wise stacking of the stable units. Coming to the influence of Ac-BIM-acid on Stm1_N^1-113^ fibrillation, it seems like 1 mM and 2.5 mM Ac-BIM-acid don’t significantly affect the amyloid-like morphology at 0 h. It starts hindering the fibril growth at 24 h, especially in the presence of the 2.5 mM Ac-BIM-acid (**Figures 3 & 4, S4-S5**). At 24 h, the fibril height seems to be unaffected while the length of the fibril is shortened in the presence of 1 mM Ac-BIM-acid (**Figures 3(E, F &H) and S4(E, F &H)**). However, in the presence of 2.5 mM Ac-BIM-acid, both the length and height of the fibril are reduced (**Figures 3(G&H) and S4(G&H)**). At 48 h, the stable units of Stm1_N^1-113^ are seen either independently or as length-wise aggregates in the presence of 1 mM Ac-BIM-acid, whereas a complete destabilization of fibrils is seen, leaving only spherical oligomeric structures, in the presence of 2.5 mM Ac-BIM-acid (**Figures 4 & S5**). The different morphological features of Stm1_N^1-113^ fibrils seen at 0, 24, and 48 h in the presence of 1- and 2.5-mM Ac-BIM-acid provide valuable information on how the compound influences the protein aggregation process with respect to time. Based on the previous studies, one can envisage that the interaction of Ac-BIM-acid to Stm1_N^1-113^ may take place via stacking and hydrophobic interactions (Campbell et al., 2008) along with the electrostatic interaction as the compound has a carboxylate functional group.

The binding of Ac-BIM-acid with Aβ42 at the interface of two of adjacent β-sheets (**Figure 6**) as indicated by docking studies suggest that such a binding may inhibit the multimerization of the fibrils. Interestingly, the binding site identified in this study is reported to be a druggable site in an earlier investigation (Marondedze et al., 2020). This perhaps be the reason for inhibition of fibril growth seen in AFM in the presence of Ac-BIM-acid (**Figure 5)**.

These findings might be particularly useful in the perspective of many amyloid-related diseases, as amyloid aggregation is considered a key pathological step in neurodegenerative disorders such as Alzheimer’s disease (AD) (investigated in this study), Parkinson’s disease (PD), Huntington’s disease (HD) and amyotrophic lateral sclerosis (ALS) (Wells et al., 2021). The ability of this compound to hinder amyloid fibril growth may provide a means to reduce the detrimental effects of mature fibril formation and plaque deposition associated with amyloid diseases. In short, the findings reported in this study suggest that Ac-BIM-acid may act as a scaffold to develop new acridine compounds against various amyloid-related disorders.

## 5. Conclusion

This study highlights the potential of newly synthesized acridine derivative 3,3’-(acridine-4,5-diylbis(methylene))bis(1-(carboxymethyl)-benzimidazolium) dibromide (Ac-BIM-acid) to bind to amyloid-like fibrils of yeast Stm1_N^1-113^ and amyloid fibrils of human Aβ42. Ac-BIM-acid compound is shown here to inhibit the growth of the Stm1_N^1-113^ and Aβ42 fibrils in a time-dependent manner. This suggests that Ac-BIM-acid can be used in blocking amyloid fibrillation.

## Acknowledgement

The authors sincerely thank Mr. Chengappa Thumisi, IIT Hyderabad, for assisting in AFM data collection. We also thank Mr. Md. Samiuddin, IIT Hyderabad, and the National High-Field NMR facility, TIFR Hyderabad, for the NMR data. PPU and Dinesh thank CSIR, and UD thanks UGC for fellowships. TR thanks SERB (Government of India) for financial support (Grant number: CRG/2022/001825). GP acknowledges SERB-CRG/2022/000714, New Delhi, India, and SOCH-IITH for financial support.

## Author contribution

TR designed and supervised the project. PPU, UD and RK carried out protein synthesis. DH synthesized the compound. KR carried out and analyzed 2D-NMR experiments. PPU and UD performed biophysical experiments and analyzed the results. AKB carried out docking studies. PPU, TR, DH, and GP wrote the original manuscript.

## Notes

### Competing Interest Statement

The authors have declared no competing interest.

### Summary of Updates

Figure 6 has been revised, as the docking data has been updated

