## Supplementary figures for "Modulation of *Saccharomyces cerevisiae* Stm1_N^1-113^and human Aβ42 amyloid fibril morphology by 3,3’-(acridine-4,5-diylbis(methylene))bis(1-(carboxymethyl)-1H-benzimidazolium) dibromide"

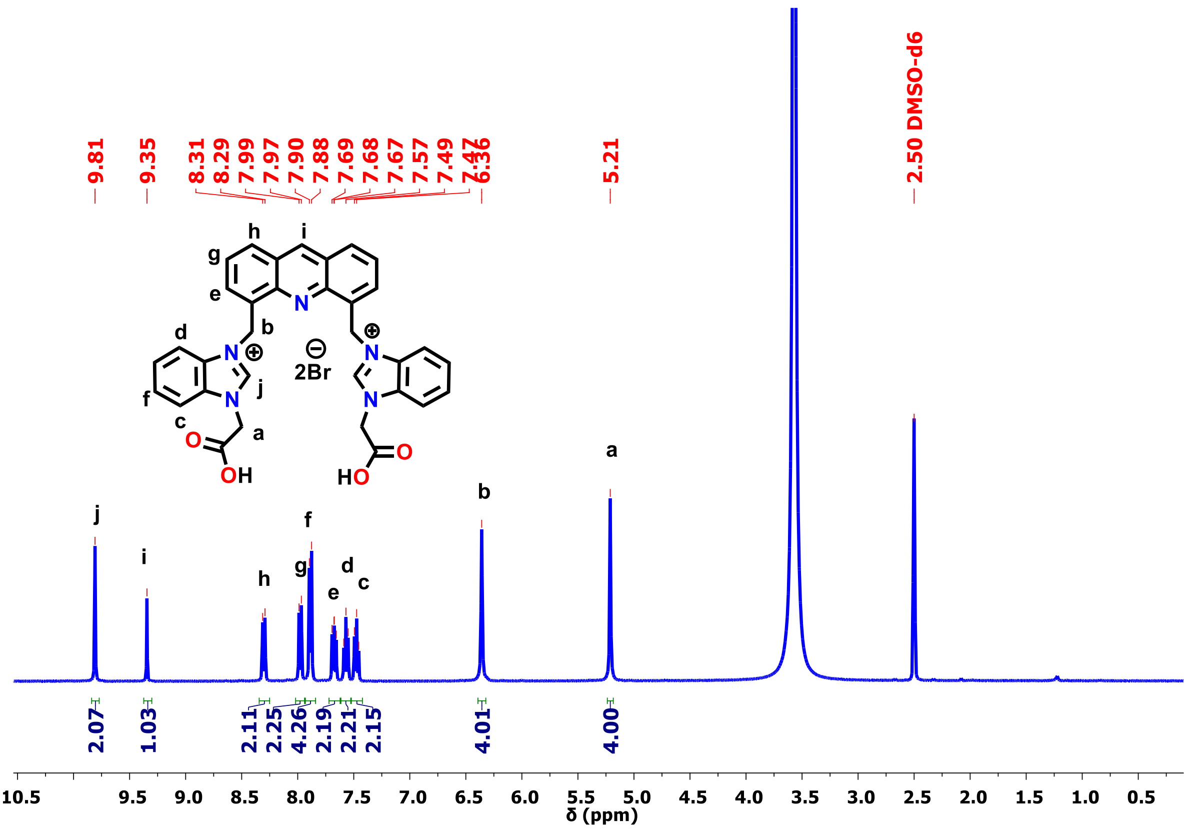


**Figure S1.** $^1$H NMR spectrum of **Ac-Bim-acid** in DMSO-$d_6$ at 25 °C

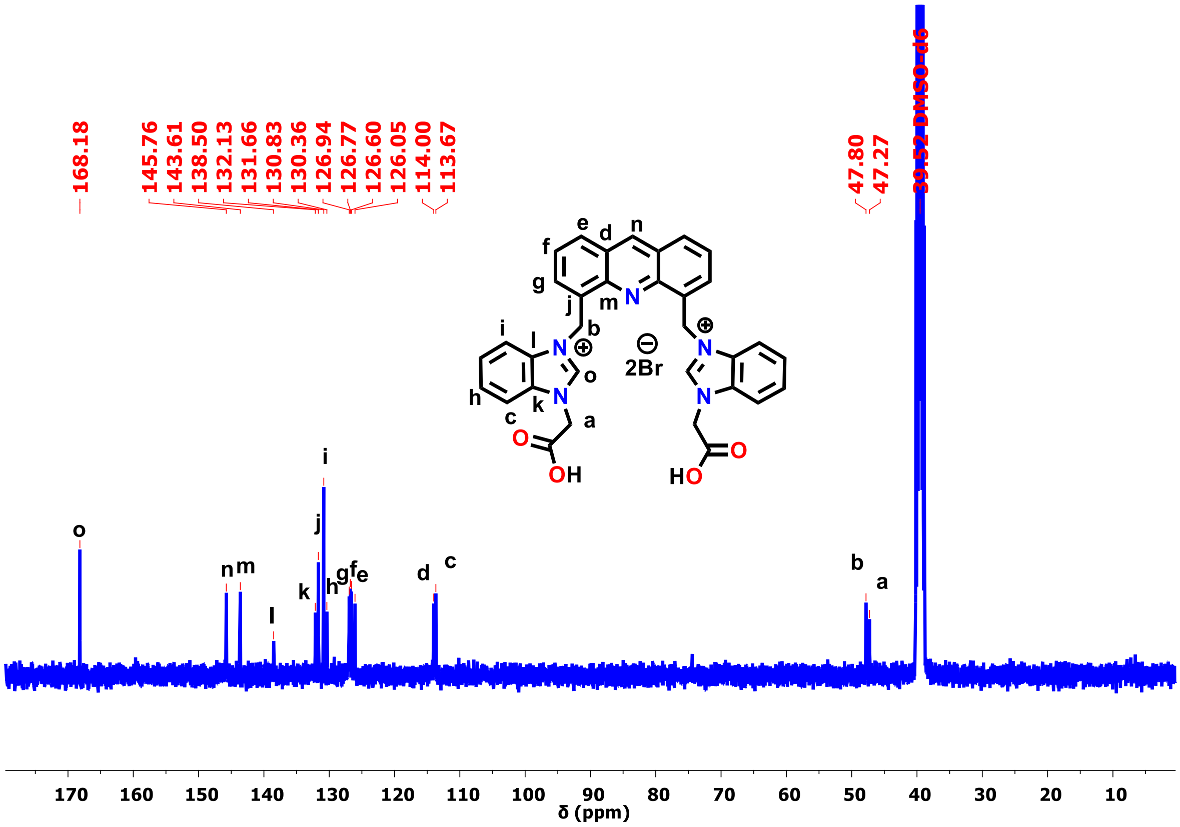


**Figure S2.** $^{13}C$ NMR spectrum of **Ac-Bim-acid** in DMSO-$d_6$ at 25 °C

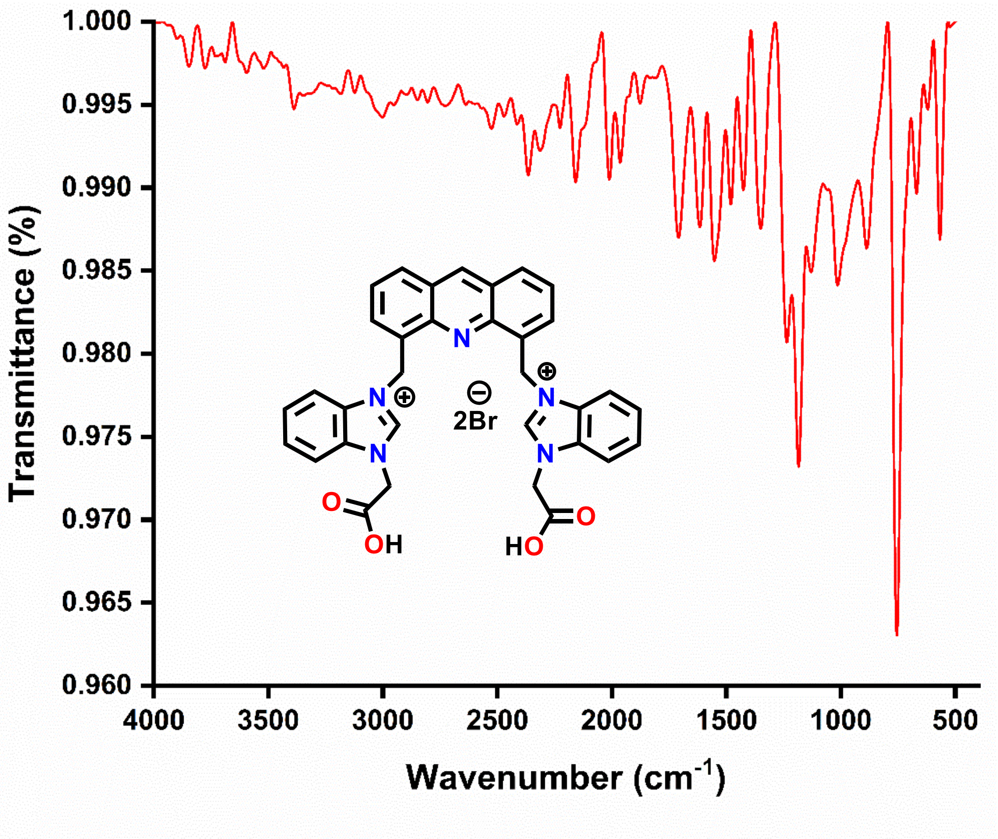


**Figure S3.** FT-IR spectrum of **Ac-Bim-acid** at 25 °C

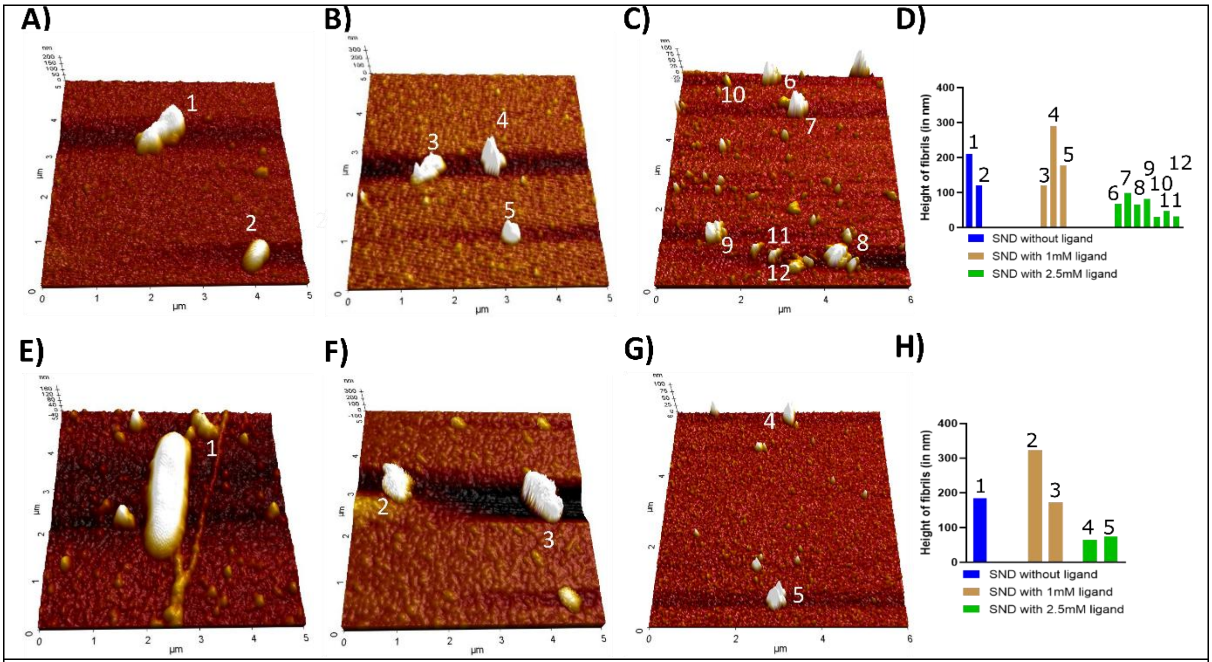


**Figure S4.** 3D-AFM images of 400 μM Stm1_$N^{1-113}$ in the absence of **Ac-Bim-acid** at (A) 0 h and (E) 24 h and, in the presence of 1 mM (B & F) and 2.5 mM (C & G) **Ac-Bim-acid** at (B) 0 h and (D 24 h. The bar plots represent the comparative height of the fibrils in the presence or absence of Ac-BIM-acid at 48 h (D & H).

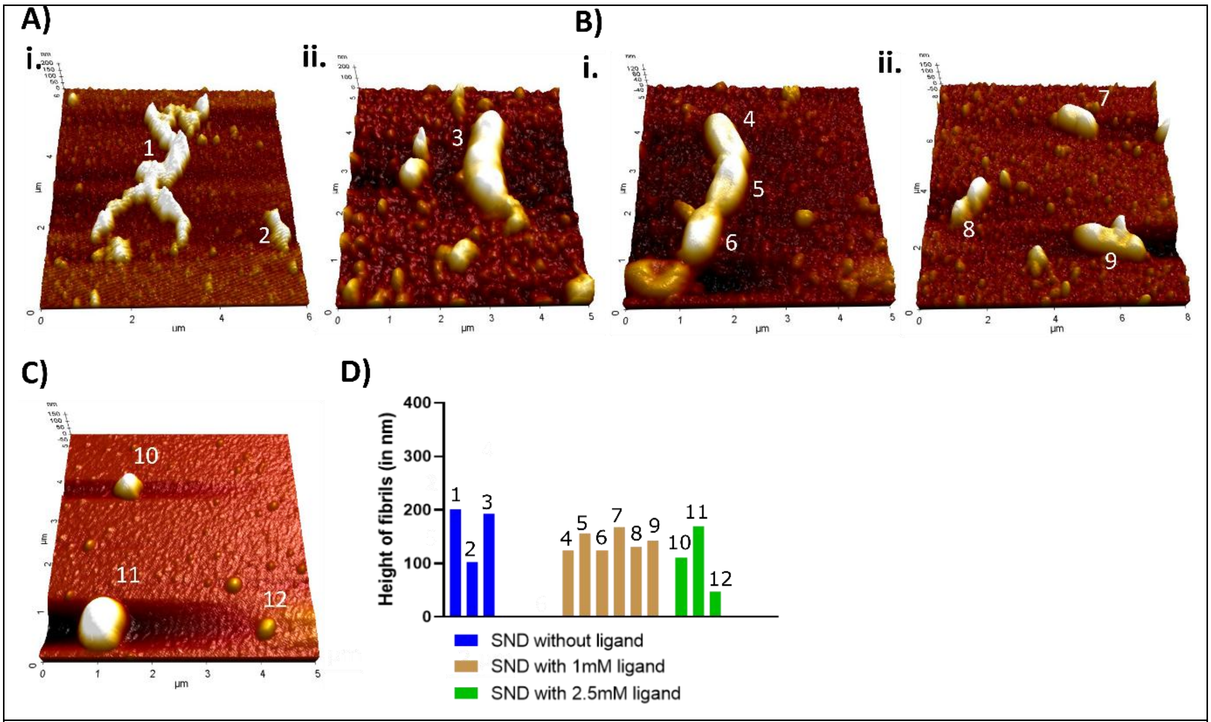


**Figure S5**. 3D-AFM images showing 2 different morphologies of 400 μM Stm1_$N^{1-113}$ fibrils in the absence of **Ac-Bim-acid** at 48 h (A) and, in the presence of 1 mM (B) and 2.5 mM (C) **Ac-Bim-acid** at 48 h. The bar plots represent the comparative height of the fibrils in the presence or absence of Ac-BIM-acid at 48 h (D).

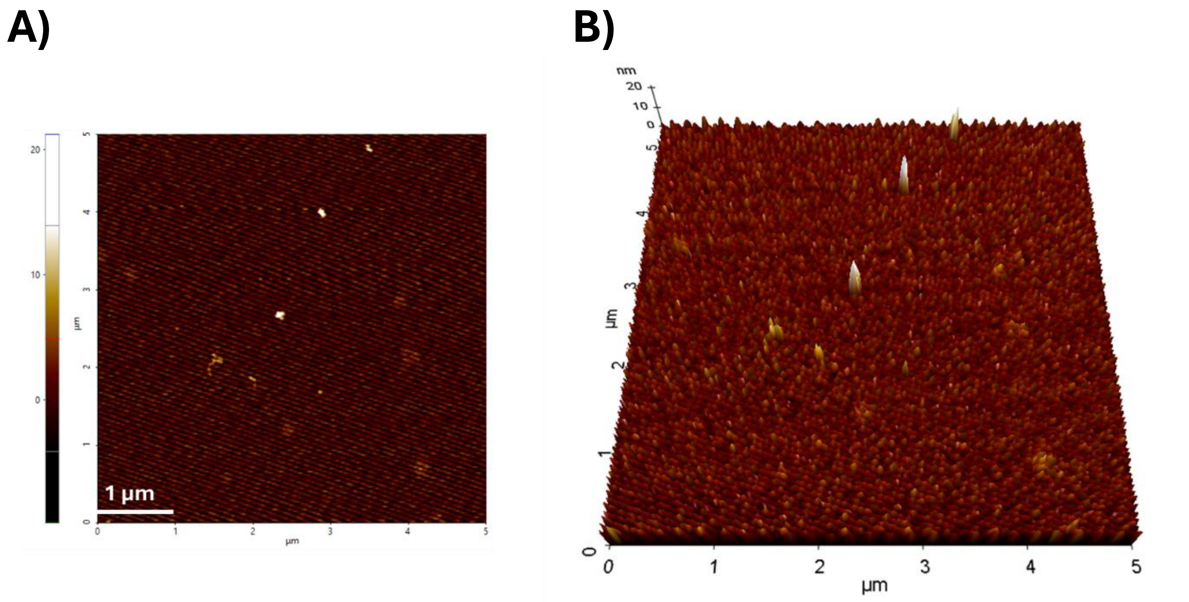


**Figure S6.** 2D (A) and 3D (B) AFM images of 1 mM **Ac-Bim-acid** at physiological salt concentration.

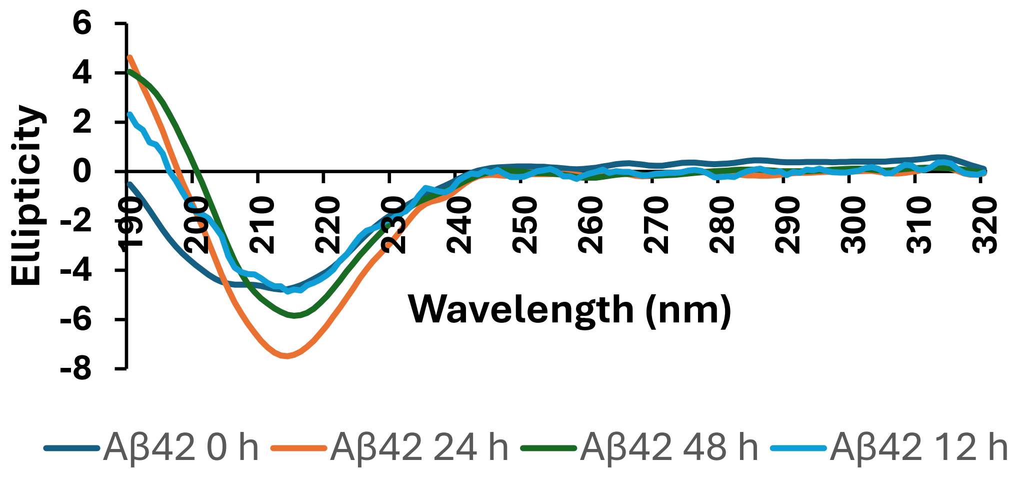


**Figure S7.** Time-dependent circular dichroism (CD) spectra of 100 µM human Aβ42 amyloid peptide showing conformational changes from 12 hours onwards.

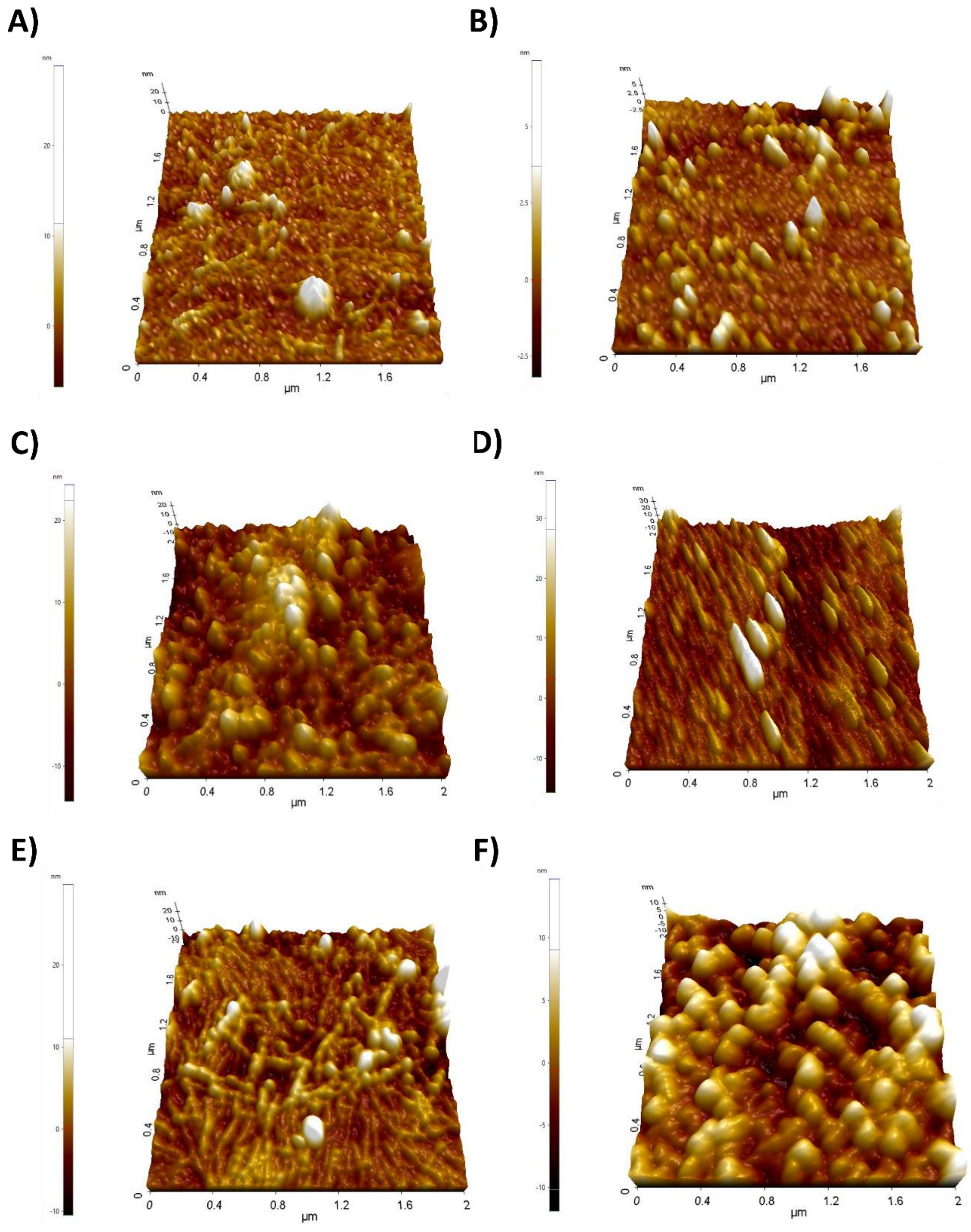


**Figure S8.** 3D AFM images of 100 μM Aβ42 at (A & B) 0 h, (C and D) 24 h and (E & F) in the absence (A, C & E) and presence of 5 mM (B, D & F) Ac-BIM-acid.
